# Accelerometer-based predictions of behaviour elucidate factors affecting the daily activity patterns of spotted hyenas

**DOI:** 10.1101/2023.05.31.543053

**Authors:** Pranav Minasandra, Frants H. Jensen, Andrew S Gersick, Kay E Holekamp, Eli D Strauss, Ariana Strandburg-Peshkin

## Abstract

Animal activity patterns are highly variable and influenced by internal and external factors, including social processes. Quantifying activity patterns in natural settings can be challenging, as it is difficult to monitor animals over long time periods. Here, we developed and validated a machine-learning based classifier to identify behavioural states from accelerometer data of wild spotted hyenas *(Crocuta crocuta)*, social carnivores that live in large fission-fusion societies. By combining this classifier with continuous collar-based accelerometer data from five hyenas, we generated a complete record of activity patterns over more than one month. We used these continuous behavioural sequences to investigate how past activity, individual idiosyncrasies, and social synchronisation influence hyena activity patterns. We found that hyenas exhibit characteristic crepuscular-nocturnal daily activity patterns. Time spent active was independent of activity level on previous days, suggesting that hyenas do not show activity compensation. We also found limited evidence for an effect of individual identity on activity, and showed that pairs of hyenas who spent more time together synchronised their activity patterns. This study sheds light on the patterns and drivers of activity in spotted hyena societies, and also provides a useful tool for quantifying behavioural sequences from accelerometer data.

## 1 Introduction

Activity patterns in animals are one of a large variety of daily rhythms such as body temperature and sleep. Activity patterns and other rhythms are governed by a variety of factors [reviewed, e.g., in 1, 2], which are not understood holistically [3]. By and large, these factors can be categorised as internal (e.g., age [4] and sex [5]), social (discussed further below), and environmental (predators and prey [6, 7] and temperature-related [8]). Daily rhythms can have a strong effect on animal survival [9–11] and reproduction [12, 13]. Yet despite their importance, field studies on animal daily rhythms and activity patterns are rare due to the difficulty of obtaining long-term data from animals whose rhythms are affected by variables of interest.

Spotted hyenas (*Crocuta crocuta*, see images in Table 1) are social carnivores found in sub-Saharan Africa. Spotted hyenas (henceforth ‘hyenas’) are well-studied with regard to their behaviour and ecology [14]. Hyenas live in large clans which can number more than a hundred individuals [15], but are typically widely dispersed and split into fission-fusion groups [16], wherein individuals and sub-groups frequently separate from each other for extended periods. Because of this fission-fusion pattern, hyenas spend varying amounts of time in close proximity with other group members [17]. Prior work on hyena activity patterns has identified considerable variability across different locations, with a common pattern being nocturnal activity with some social behaviours peaking during twilight hours [18, 19]. During the day, they typically spend their time at rest, sometimes in and around communal dens, while at night they typically hunt and scavenge for prey, often with other members of their clans.

**Table 1:**
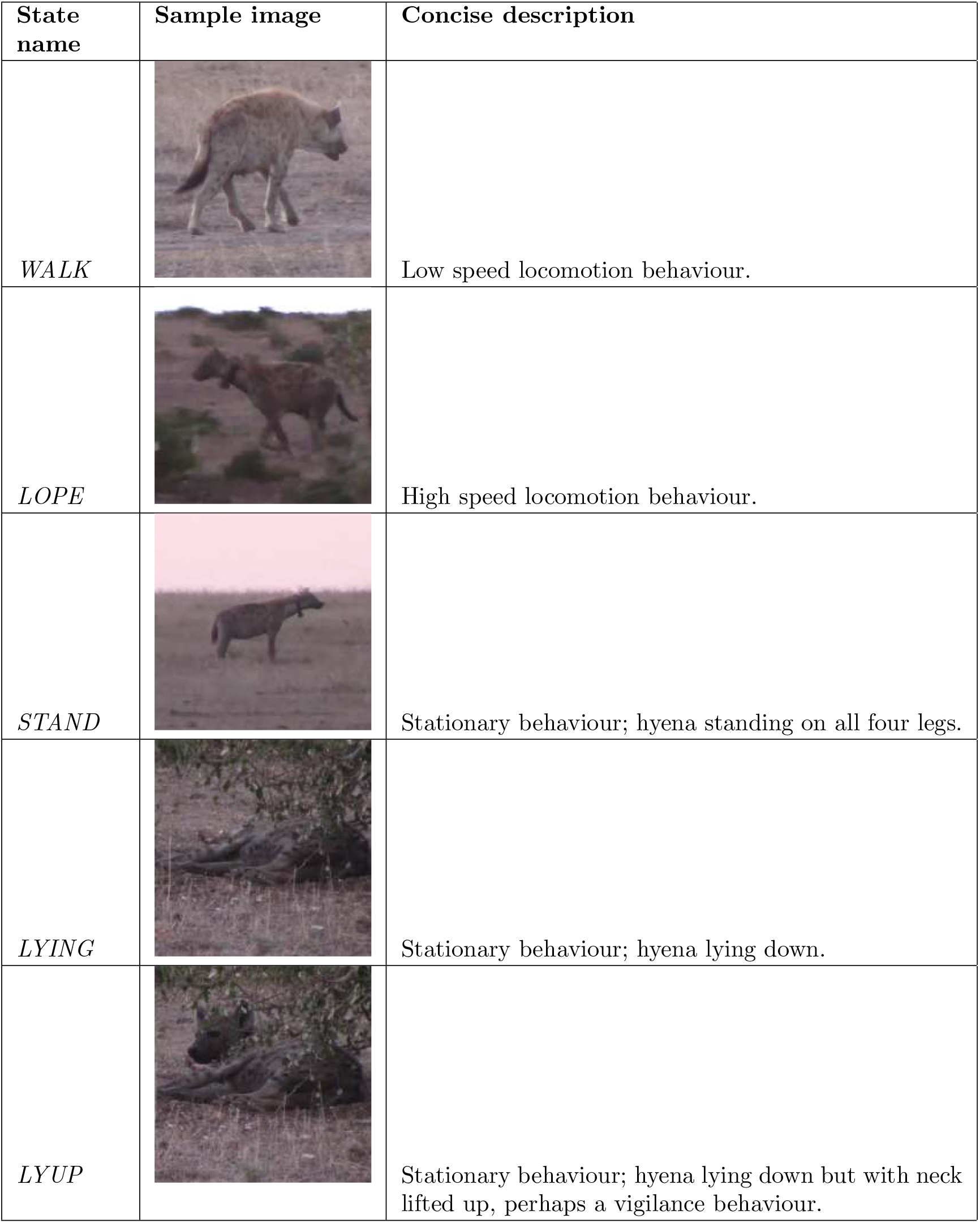
Hyena behaviours considered for this analysis, with brief descriptions. Images are from video data used for noting ground-truth, used in training our classifier (see Methods).

| State name | Sample image | Concise description |
| --- | --- | --- |
| *WALK* |  | Low speed locomotion behaviour. |
| *LOPE* |  | High speed locomotion behaviour. |
| *STAND* |  | Stationary behaviour; hyena standing on all four legs. |
| *LYING* |  | Stationary behaviour; hyena lying down. |
| *LYUP* |  | Stationary behaviour; hyena lying down but with neck lifted up, perhaps a vigilance behaviour. |

For group-living species such as hyenas, social interactions may be an important driver of activity patterns. For example, individuals may stimulate one another to transition from a resting behaviour to a more active one, or individuals may prefer to rest together simultaneously either for safety or social bonding. Social entrainment occurs when cyclical activity patterns or circadian rhythms become aligned across individuals. While most previous literature does not explicitly address the social entrainment of daily activity patterns, the social entrainment of *circadian rhythms*, more broadly, has been analysed to a larger extent [e.g., 20–22]. While social entrainment is likely to be a widespread driver of activity patterns, its study is hampered due to two main challenges. The first problem arises because it is fundamentally very difficult to quantify the role of inter-individual entrainment as opposed to entrainment by a common temporally changing source (such as sunlight or temperature). The second problem arises because the study of within- and among-individual variation in circadian rhythms requires recording activity of the same individuals repeatedly over the day and night. In contrast, population or species level estimates of activity patterns are productively studied in wild animals using camera traps [23]. The challenges associated with studying activity patterns at the level of individuals are often addressed using experimental studies on pairs of captive animals [reviewed in 24], where the environment can be carefully controlled and individuals monitored continuously. While experimental work with captive animals does provide tremendous insight into the mechanics and physiology of the question of social entrainment, addressing the ecological and behavioural aspects of this question also requires long-term observational studies of animals in their natural habitats.

Recent advances in bio-logging technology [25] open up new possibilities for the continuous monitoring of animal behaviour in the wild, allowing us to begin tackling these topics in natural settings. In particular, accelerometer loggers deployed on animals offer the potential to capture behaviours remotely at a fine scale and over long periods of time. A tri-axial accelerometer is a device that measures acceleration in three dimensions (front-back or *surge*, up-down or *heave*, and sideways or *sway*). Using tri-axial accelerometers in combination with machine-learning approaches [e.g., 26–31], we can predict the behavioural states of animals for the period of deployment with a high temporal resolution, allowing us to obtain long-term observational data on individual animals.

Such long-term behavioural sequences offer the opportunity to address both long-standing and new questions about the drivers of activity patterns in the wild. For instance, we can test whether animals compensate for days of high activity on the following days. This question is motivated by a similar phenomenon documented in sleep [32], known as sleep debt, where individuals make up for lost sleep. Sleep is, however, different from activity, since active individuals are always awake, whereas waking individuals can be either active or inactive. The question is thus whether hyenas make up for days of high activity by being less active on subsequent days, and vice versa. While there is some evidence both for [33] and against [34] activity compensation in humans, asking this question about non-human animals in the wild has been difficult so far. Additionally, we can test whether animals in nature have significantly idiosyncratic (individual-specific) activity patterns, a question so far addressed mainly in humans [35] and lab animals [2] (but see [36, 37]). Once again, we can draw parallels to research on sleep, where consistent variation among individuals is seen in their preferred times of sleeping and awakening, also referred to as an individual’s chronotype [38, 39]. Furthermore, individual animals are known to be more or less active in certain contexts depending on their personality [40]. Finally, to address potential social drivers of activity patterns, we can test whether animals in the same social group tend to synchronise their activity patterns.

Here, we combine accelerometer data with machine learning models to characterise activity patterns in spotted hyenas. We develop and validate a random forest classifier that can predict hyena behavioural states from accelerometer data, using video observations of hyenas as ground truth. We further confirm the validity of our approach by comparison to previous studies. We then investigate the role of three potential and non-exclusive factors that could affect hyena activity patterns: activity compensation between consecutive days, individual idiosyncrasies, and social synchronisation of activity patterns. We test three hypotheses with regard to activity patterns: 1. that high activity on a day will imply lower activity on the next day (the activity compensation hypothesis); 2. that individual hyenas’ activity patterns are idiosyncratic, and the identity of an individual greatly influences its activity pattern (the individual idiosyncrasies hypothesis); and 3. that some pairs of hyenas (specifically, those that spend time in close proximity) have synchronised activity patterns (the social synchronisation hypothesis).

## 2 Methods

### 2.1 Data collection and pre-processing

Data were collected using tracking collars deployed on five adult female spotted hyenas (named WRTH, BORA, BYTE, MGTA, FAY; same names used here throughout) who were members of the same clan, located in the Masai Mara National Reserve, Kenya. Hyenas were part of a long term individual-based study with access to information on genealogical, demographic, and social relationships among individuals [41]. Collars contained a custom-designed tag recording audio data at 32 kHz, tri-axial accelerometer data at 1000 Hz, and magnetometer data at 1000 Hz (DTAG; Mark Johnson). This tag was integrated with a GPS logger recording at 1 Hz (Gipsy 5; Technosmart, Accuracy = 95% of points in ¡ 5 m) which also provided time synchrony accurate to the nearest second. The integrated tag was then housed in a Tellus Medium collar containing a VHF transmitter, additional (low-resolution) GPS module, Iridium transmitter/receiver, and automatic drop-off unit (Followit Sweden AB). Collars were deployed in December 2016, began recording simultaneously on January 1st, 2017, and were in operation continuously for approximately 40 days (Table 2). Available triaxial accelerometry data were down-sampled to 25 Hz for further analysis. GPS data were filtered to exclude unrealistically high velocities for individuals (thresholding at the 99.95th percentile *≈* 14.8 ms^*−*1^). Missing GPS sequences of short duration (*<* 5 s) or short displacement (*<* 5 m) were replaced by linear interpolations.

**Table 2:**
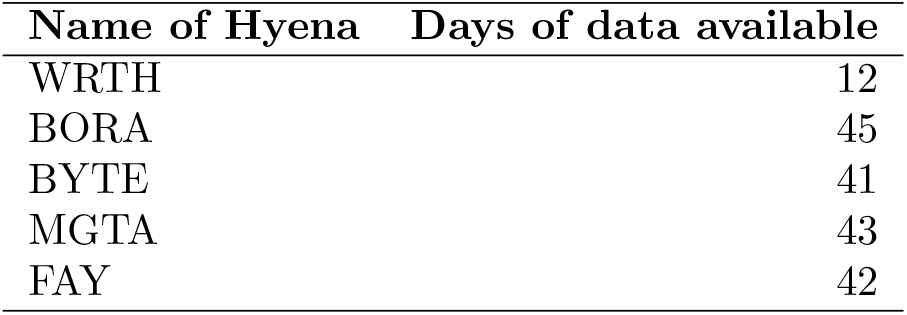
Summary of available data.

Ground-truth behavioural state recordings were used to train the behavioural classifier. To do so, we opportunistically used videos recorded during playback experiments of long-distance recruitment (“whoop”) calls [42] to hyenas wearing collars in the field. During these experiments, a sequence of calls was played to initially-resting hyenas. These experiments often catalysed strong responses, where hyenas went from resting to eventually walking or loping in the direction of the calls, thus exhibiting a range of basic, movement-based behavioural states. 2-4 separate videos were collected for each of four tagged hyenas (data for the individual WRTH could not be used since that collar failed on day 13, see Table 2), such that the total video coverage of each was approximately one hour. These videos were synchronised with DTAG data and then manually transcribed to behavioural state text files (hereafter referred to as ‘audits’), and simplified to a five-behavioural-states model (also called an *ethogram*, Table 1).

### 2.2 VeDBA-based activity levels

All analyses were carried out in the Python 3.10.6 [43] programming language. To obtain preliminary behavioural labels, we used a simple, widely used metric based on activity: the Vectorial Dynamic Body Acceleration (VeDBA: [44]). VeDBA is a radial metric, proportional to activity output, and independent of the direction of the acceleration vector. Using a sliding 1 second time window over the data, the VeDBA was computed by 1. finding the dynamic acceleration (acceleration above the mean for the time window) at each point, and 2. adding the norms of the dynamic acceleration vectors across the time window. We then divided the available data into non-overlapping 3-second intervals each containing 75 values of VeDBA. For each interval, we calculated the log mean VeDBA. We then visualised the distributions of log mean VeDBA values separately for each individual, as well as for the aggregate dataset. Separate peaks seen in this distribution were used to categorise VeDBA-based activity levels. These categories, the VeDBA-based activity levels, were defined based on local minima in of the distribution of log mean VeDBA.

### 2.3 Classifier design

Our goal was to identify behavioural classes from our ethogram for every 3-second interval spanned by the collected data. To do so, we first performed feature extraction using 16 features (*{*min, max, mean, and variance} of {surge, sway, heave, and VeDBA} values for the 3-second interval). These features from all audits were used to train three classifiers: a random forest (RF), a support vector machine (SVM), and a *k* -nearest-neighbours (*k* -NN) classifier. These classifiers were trained and tested using the Python package scikit-learn 1.2.2 [45], and were initialised using the default parameters in this package.

To validate the performance of these classifiers, we used three methods. In the first method, the classifiers were trained with a random 85% of the data, and then tested on the remaining 15%. This method is simple, commonly used, and provides basic technical information about the classifiers’ performance. However, the scores generated using this method can show spuriously high accuracy values, because of over-fitting within the same behavioural audits. These inflated values may occur because 3-second windows within a single recording are not really independent of one another (for instance, consecutive 3-second intervals are most likely of the same behavioural state). Our second method of performance estimation was therefore to train the classifiers on subsets of the data, each time leaving one audit out, training on the remaining audits, and then testing on the audit that was left out. This audit-wise approach ensures greater independence between test and training data. Finally, to address the question of generalisability across individuals, our third approach was to test the performance of our classifier on the data of each hyena, after training the classifiers on data from the rest of the hyenas. For each method, we computed the overall accuracy and generated a confusion matrix.

Additionally, to test how well a simple VeDBA-based characterisation captured hyena behaviours, we looked at the behavioural state composition of each VeDBA-based activity level (from 2.2).

### 2.4 Quantifying hyena daily activity patterns

The classifier with the highest performance, the Random Forest classifier (See Results), was trained with all available ground-truth, and then made to predict behavioural states across the entire period of collar deployment for each hyena. These predictions were used to infer the hyenas’ daily activity patterns. For each hyena, we calculated the fraction of time spent in the active states (*WALK* and *LOPE*) for each hour of the day.

### 2.5 Activity compensation across consecutive days

To test the activity compensation hypothesis, for each hyena, we plotted the activity on day *i* + 1 against that on day *i*. We then performed linear regressions for these data for each hyena. The activity compensation hypothesis predicts a negative correlation between activity patterns on consecutive days.

Because it is possible that activity compensation occurs on a timescale longer than a day, we also explored whether high activity during the past *m* previous days was associated with lower activity on a given day. To do so, we found the average activity between day *i* − *m* + 1 and day *i*, and performed a regression of this average activity with activity on day *i* + 1. To avoid introducing too many comparisons which could result in spurious relationships, we restricted this analysis to *m* = 2 and *m* = 5 days.

### 2.6 Idiosyncratic activity patterns

To test whether the daily activity patterns of different individuals differed consistently from one another, we compared activity levels in each hour of the day both within and across different hyenas.

We first computed *activity curves* for each hyena on each day, here defined as the percentage of time an individual was in an “active” state (*WALK* or *LOPE* as predicted by our classifier) during each hour of the day. The activity curve for each day for each hyena was represented as a 24-component vector of percentages. We then assessed the variability between pairs of different days by computing the sum of squared differences between these vectors. For within-individual variability, we compared activity curves from the same hyena on different days, whereas for across-individual variability we used activity curves from different hyenas on different days. To quantify the difference between within- and across-individual variability, we defined the individuality statistic as the difference between the mean across-individual and the mean within-individual variabilities (see Appendix B for more details).

We next used a permutation test to determine whether the within-individual variability was significantly lower than the across-individual variability. For each permutation, we created 5 pseudo-hyenas by randomly shuffling activity curves amongst individuals on each day, such that the pseudo-hyenas had the same amounts of data as the real ones (Table 2), then computed the value of the test-statistic defined above. Using 5000 permutations of the data, we defined a null distribution of the test statistic. Finally, we tested for a significant difference using the conventional definition of the *p*-value (one-tailed, since we only test whether the within-individual variability is less than across-individual variability), with a significance threshold of 0.05.

### 2.7 Synchronisation of activity patterns and relation to proximity

We tested whether pairs of hyenas tended to be more synchronous in their activity patterns than expected by chance. To quantify synchronisation of daily activity patterns between pairs of hyenas, we first computed the series of hourly activity for each hyena from the beginning to the end of the deployment. For each pair of individuals, we then computed a synchronisation score for their activity patterns (Appendix B). After computing these scores, we randomly shuffled each hyena’s sequence of 24-hour activity curves (i.e, the order of days was shuffled, while the order of hourly activity levels in each day was preserved). We then recomputed these scores with the permuted data. This permutation was repeated 100 times in each case, letting us define a ‘null hypothesis’ that captures the distribution of the synchronisation score for a pair of hyenas expected purely by chance. A pair of hyenas was said to be synchronised overall if its true synchronisation score was greater than 95% of the scores generated by these permutations.

To test the role of proximity in synchrony, for each unique pair of hyenas, we computed the proportion of available data wherein they were within 200 m of each other (the standard distance threshold used by the long-term field study to define individuals as part of the same subgroup [46]). We then defined a network of proximity where edges were defined as the proportion of time two individuals were within 200 m of one another. Given our limited sample size of only 10 unique pairs of hyenas in this study 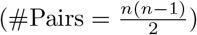, we chose to compare the networks of proximity and synchronisation qualitatively rather than performing more formal statistical analyses. We also repeated the above analysis with thresholds 50 m, 100 m, 200 m, 300 m, and 500 m to test for robustness.

## 3 Results

### 3.1 Three VeDBA-activity levels

Across all hyenas in our study, the log mean VeDBA was distributed with three distinct peaks (Figure 1) that occurred at similar values. This trimodal distribution suggests that hyenas display three basic activity levels (*low, medium*, and *high*), which provides a rough, preliminary description of their behaviours.

**Figure 1:**
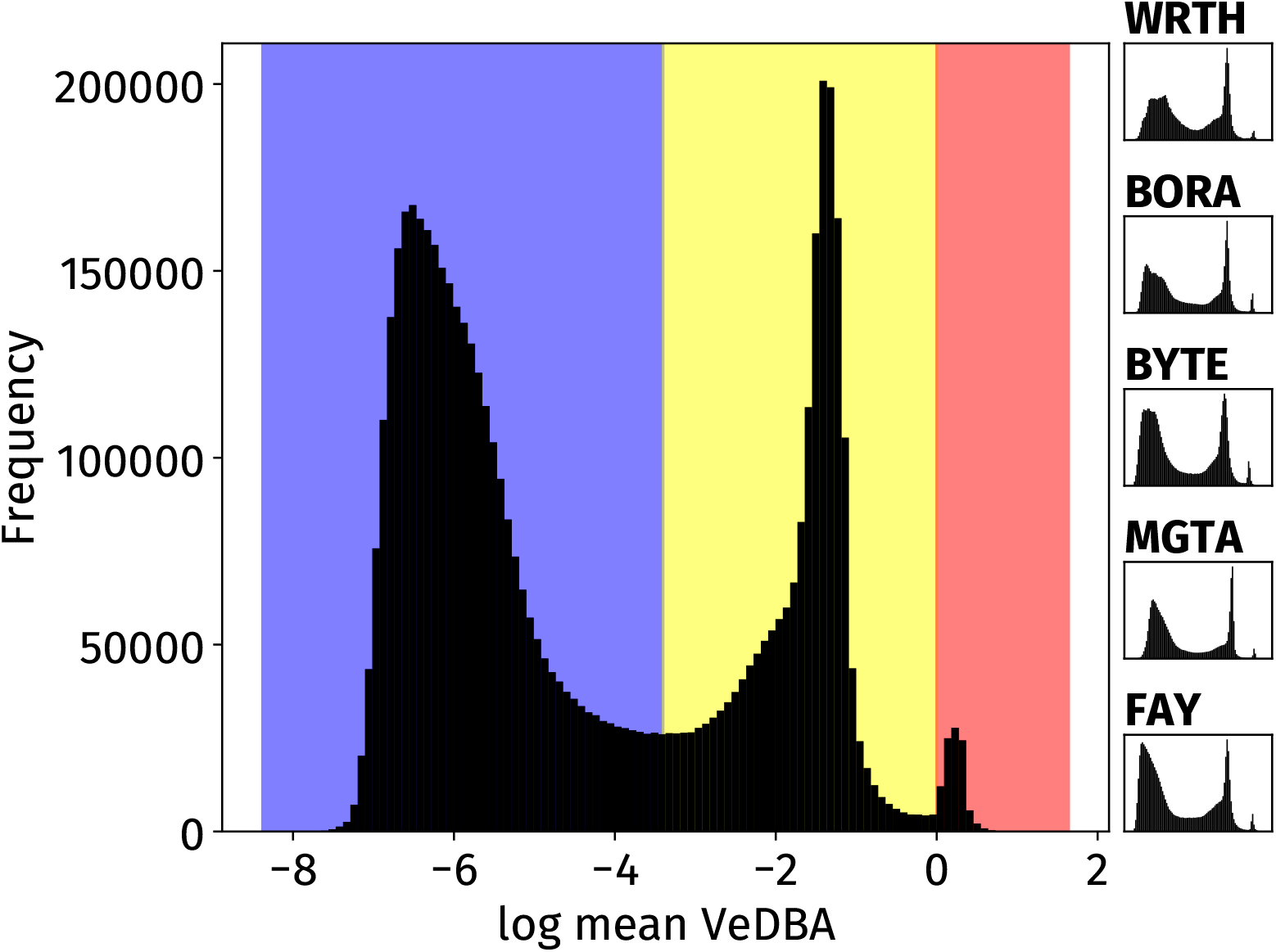
Histograms of log mean VeDBA values. Averaging was across 3 s intervals. 3 separate peaks are seen, corresponding to three levels of activity, labelled *low* (*x* < − 3.4), *medium* (− 3.4 < *x* < 0), and *high* (*x* > 0). These activity levels are shown respectively coloured blue, yellow, and red. The same 3-peak pattern was observed separately in all hyenas (insets at right).

### 3.2 Classifier performs well with spotted hyena accelerometry

Across all classifiers, the Random Forest classifier performed best and is therefore presented here (for results of other classifiers, which performed similarly well, see Appendix A). The random forest classifier had a 92% accuracy in the randomised testing on 15% of the data, and performed well for all behaviours (Figure 2). When tested across audits, the performance was reduced as expected but the classifier still performed reasonably well with an accuracy of 83%. Finally, testing across individuals also yielded appreciable performance (78% accuracy, suggesting at least some generalisability across individuals). In all cases, the classifier was most likely to confuse the *STAND* and *LYUP* states, which accounted for most of the misclassification.

**Figure 2:**
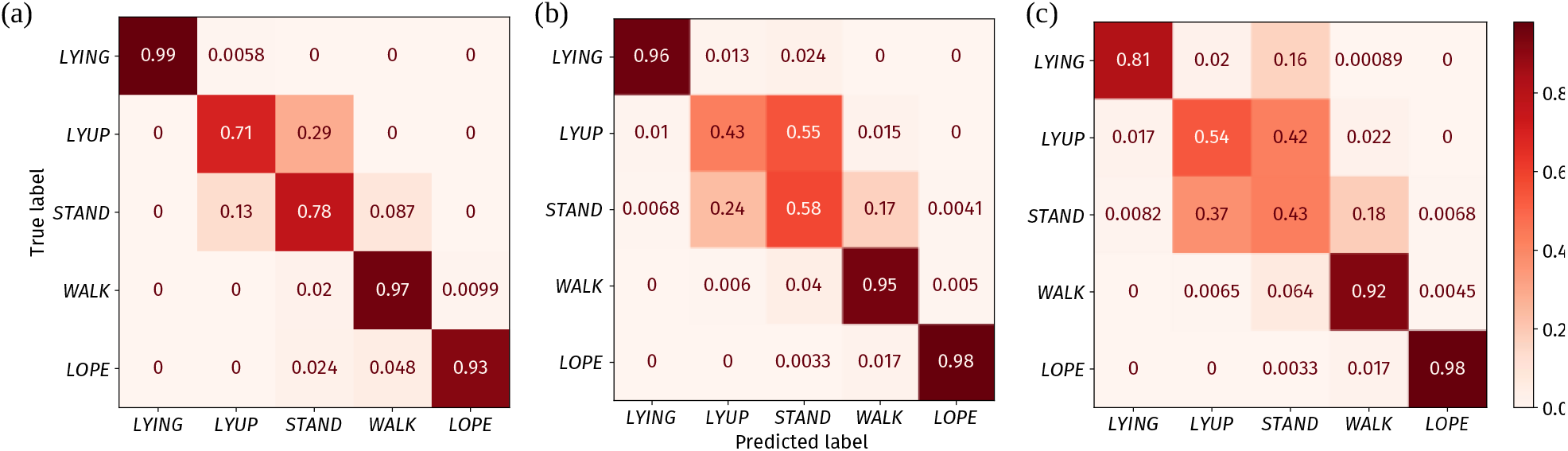
Confusion matrices for a random forest classifier trained and tested on data which were separated (a) randomly; (b) audit-wise; and (c) individual-wise (see Methods). The rows of each confusion matrix represent the true behavioural state of an animal (from the ground-truth) and the columns represent the classifier predictions. Each square *A*_*ij*_ shows the fraction of states *i* (y-axis) interpreted by the classifier as state *j* (x-axis). Higher values for the diagonal elements indicate a more accurate classifier. The classifier appeared to perform well in all cases. The *LYUP* and *STAND* behaviours were most often confused with one another.

VeDBA-based activity levels corresponded to behavioural states in an intuitive manner (Figure 3). The low activity level corresponded almost entirely to static behavioural states, and the high activity level was comprised almost exclusively of the *LOPE* state. The medium activity level was more ambiguous, and consisted mainly of the static *LYUP* and *STAND* states (which themselves were often confused by our classifier) as well as the dynamic *WALK* state. Overall, the results show that VeDBA-based classification of activity patterns can capture the broad patterns of activity, and moreover, that our classifier allows for a more detailed breakdown of activity states into behaviours.

**Figure 3:**
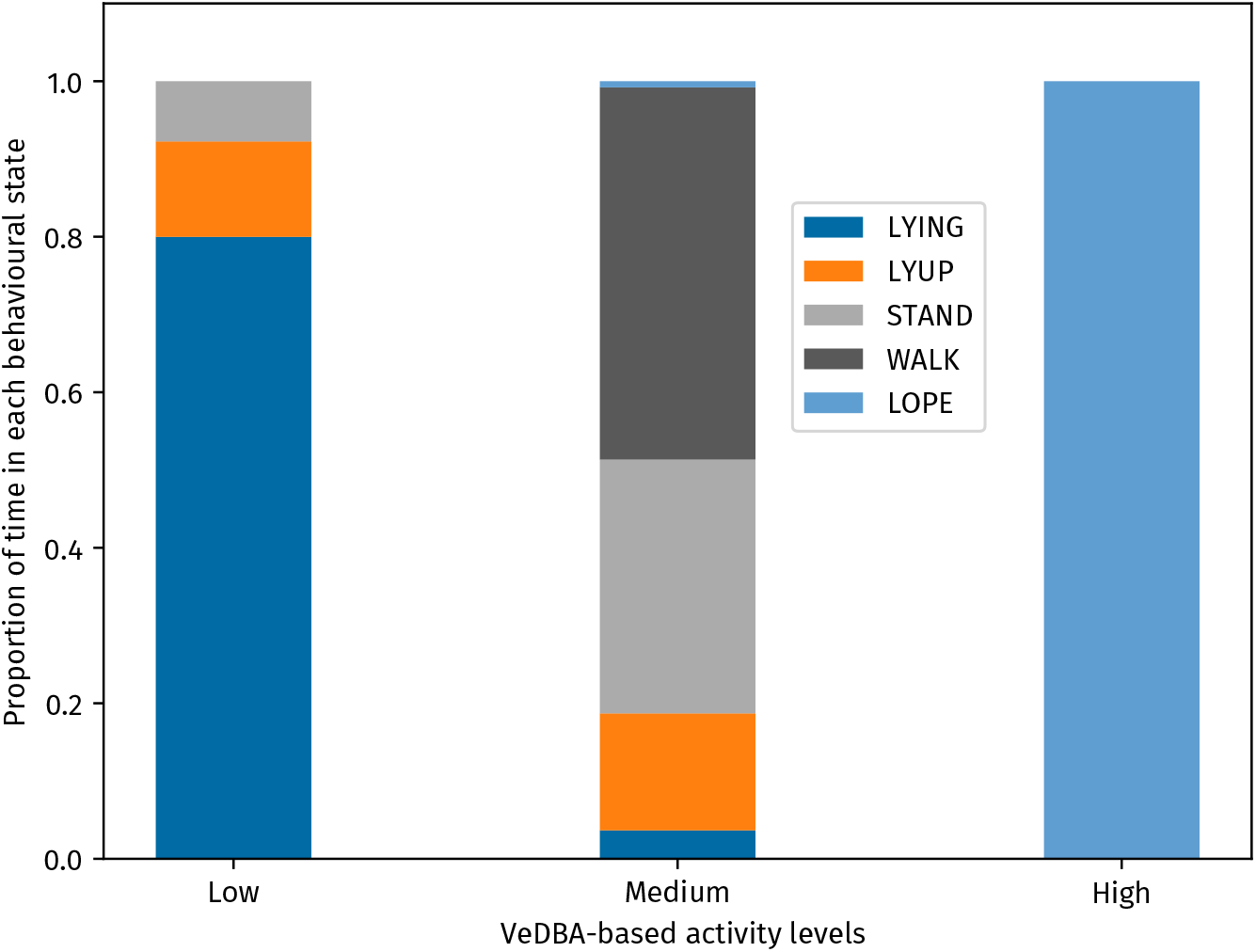
Behavioural composition of the three VeDBA-based activity levels, as revealed by the behavioural classifier. Behavioural state classifications and VeDBA-based activity levels were assigned for non-overlapping 3 s time-windows. The *high* activity level consists nearly entirely of the fast movement state *LOPE*, while the *low* activity level consists of non-movement behaviours (*LYING, LYUP*, and *STAND*). The *medium* activity level is composed mainly of the stationary behaviours *STAND* and *LYUP*, as well as the moving *WALK* state.

### 3.3 Classifier predictions display 24-hour activity pattern

Hyena daily activity patterns as predicted by our classifier indicate that all hyenas in our study were nocturnally active while largely resting through the day (Figure 4).

**Figure 4:**
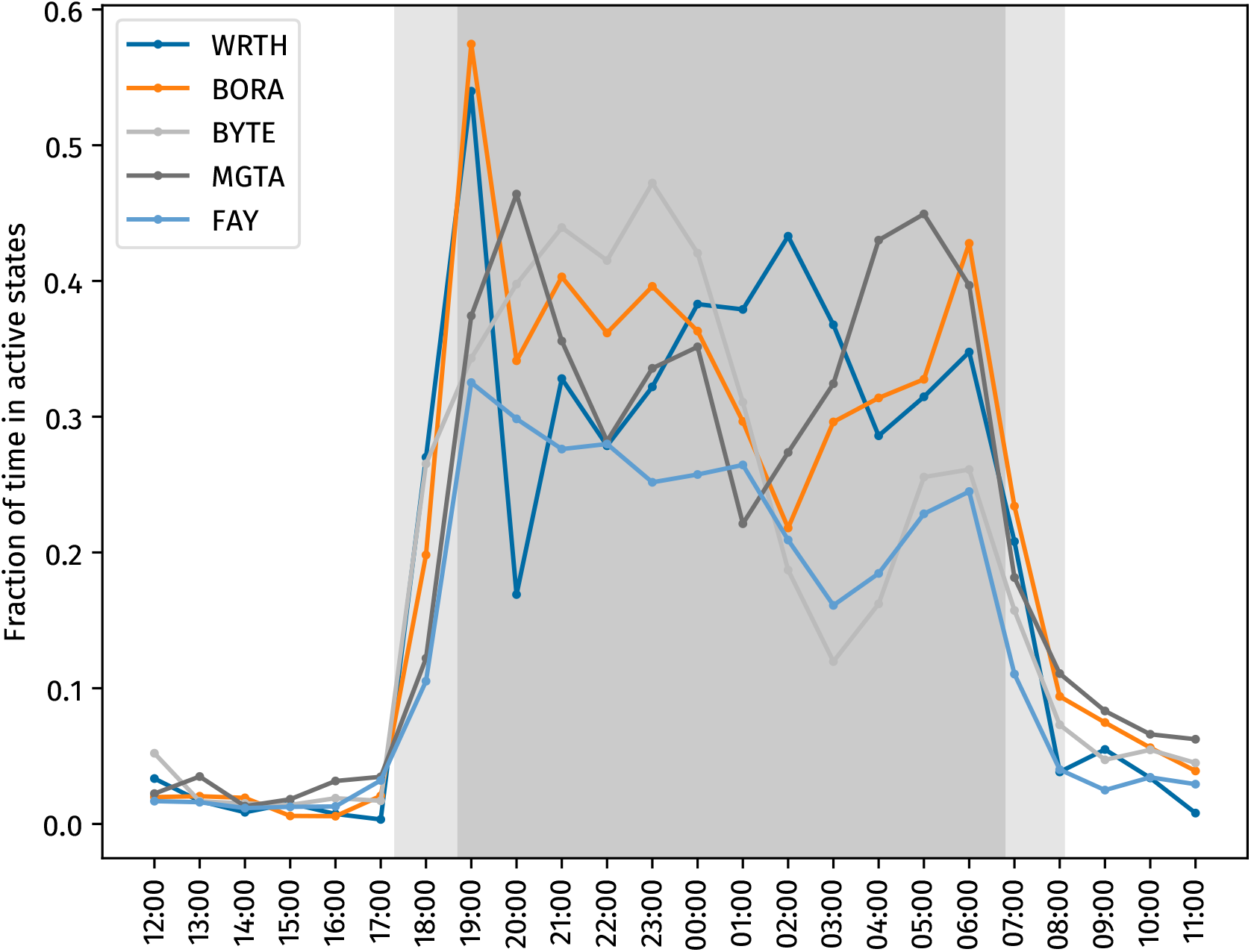
Fraction of time spent in the *WALK* or *LOPE* movement states for each hyena (coloured lines), at each time of day (x-axis). The area shaded dark grey represents night-time, and areas shaded light grey indicate twilight.

### 3.4 No activity compensation on consecutive days

Linear regressions between activity on days *i* and *i* + 1 showed no clear trends, indicating that, in the five collared individuals, activity on a given date was unrelated to activity on the next day (Figure 5). Further, no effect was seen when accounting for average activity on the preceding m = 2 or m = 5 days. (Appendix C).

**Figure 5:**
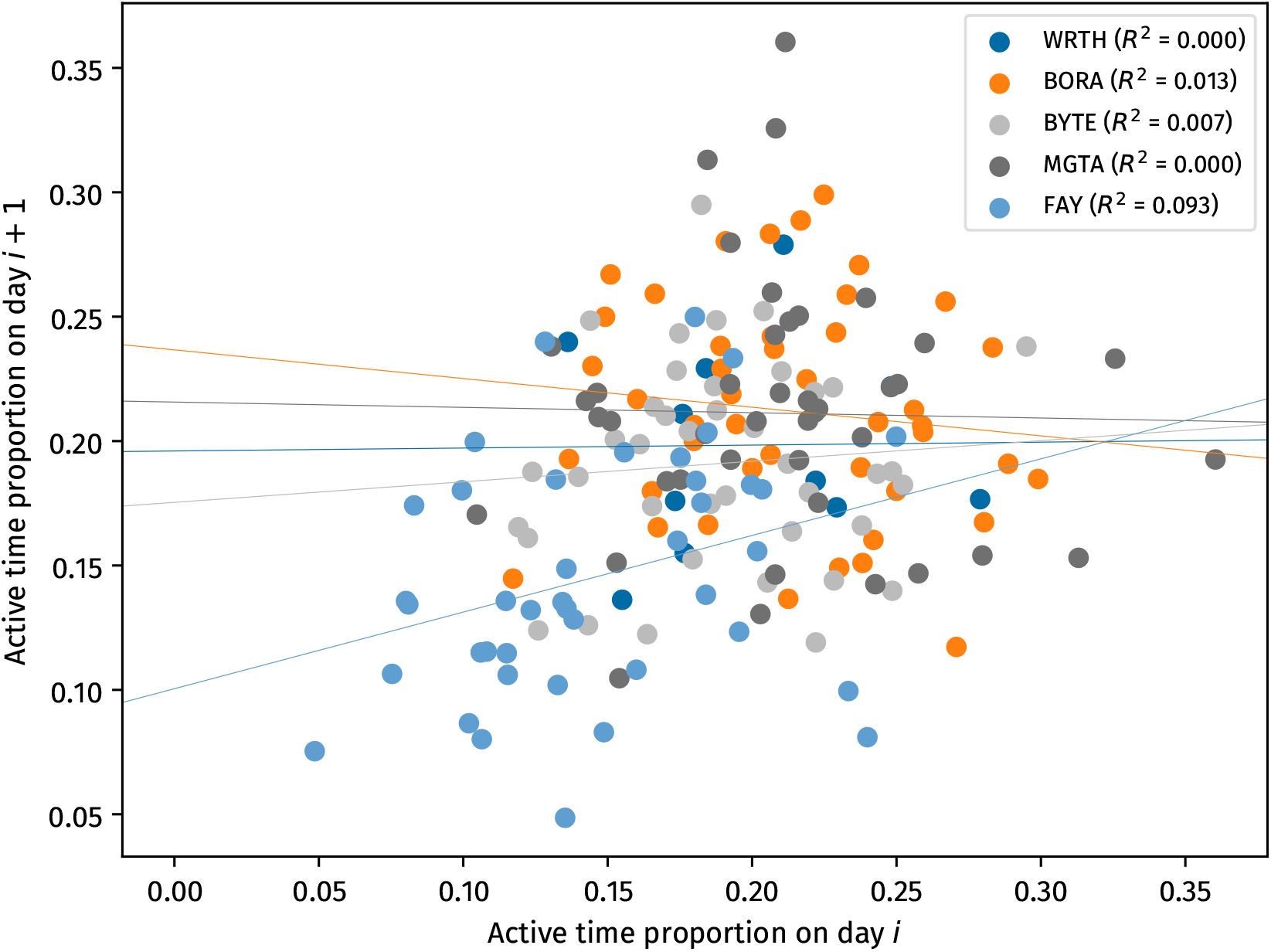
Relationship between activity levels on subsequent days for all hyenas. Each data point represents a day *i, i* + 1 pair. Different colours indicate different individuals, with lines showing linear regressions for each individual. No consistent trend is observed, suggesting that hyenas do not adjust the current day’s activity levels based on their activity on the prior day.

### 3.5 Individuals show idiosyncratic activity patterns

Within-individual differences between activity curves were lower than across-individual differences (*p <* 1/5000). However, while statistically significant, the distributions of the within- and across-individual differences had large, overlapping spreads (Figure 6). In other words, the range of variability in activity patterns within individuals across days was on balance less than that across individuals, however this difference was small and may not be practically meaningful.

**Figure 6:**
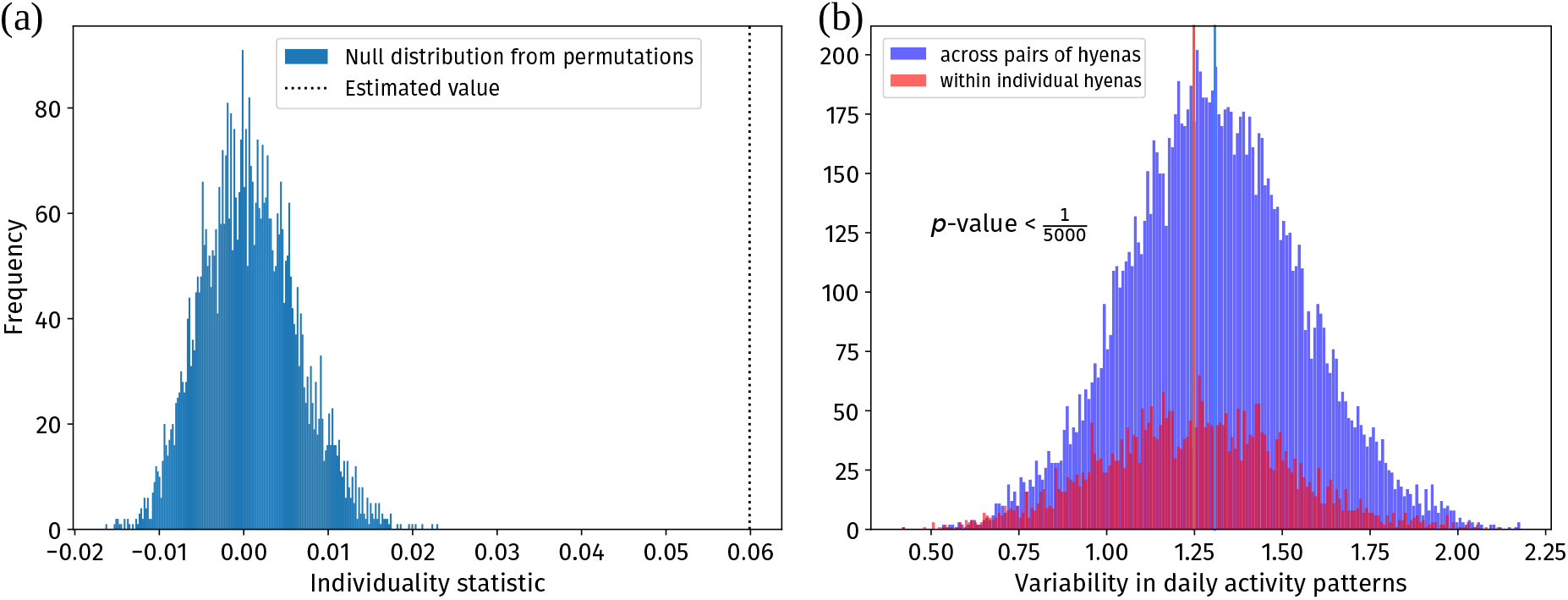
(a) Estimated value (dotted black line) of the individuality statistic exceeded those predicted to occur by chance (blue histogram), indicating a significant difference between within- and across-individual variability of daily activity patterns. (b) Histograms of within-individual (red) and across-individual (blue) variability in activity patterns. Mean values are shown as bold vertical lines of corresponding colours. Permutation tests show that the separation between the means, while statistically significant, has a small effect size as the two distributions overlap substantially.

The above pattern might be driven by all individuals showing some consistency in their activity patterns or by only some of them doing so. We therefore conducted a follow-up analysis in which we compared daily variability in a hyena’s variability in activity patterns across days to the overall variability across all individuals and all days. This analysis indicated that hyenas were slightly more individualistic than expected from chance, but this was not statistically significant. Additionally, while some individuals showed lower within-individual variability than the variability across individuals, one individual (MGTA) actually showed higher within-individual variability (Appendix D).

### 3.6 Some pairs of hyenas synchronise their activity patterns

Four out of ten unique pairs of hyenas had more synchronised activity patterns than expected by the null model permutations (Figure 7), with the rest showing levels of synchrony consistent with that expected based on daily activity patterns. In particular, BORA showed higher than expected synchrony with WRTH, BYTE, and FAY, while BYTE and FAY also showed higher than expected synchrony. Comparing to patterns of proximity revealed that lack of proximity seemed to rule out synchronisation, but synchronisation was not guaranteed when a pair of hyenas spent a large amount of time in proximity. For example, MGTA, the individual least synchronised with other hyenas, also spent the least time in proximity with any of them. In contrast, the pairs WRTH-BORA and WRTH-FAY spent substantial time in close proximity, but did not synchronise their activity patterns. The results remained consistent when we used different threshold distances for computing the proximity networks (Appendix E).

**Figure 7:**
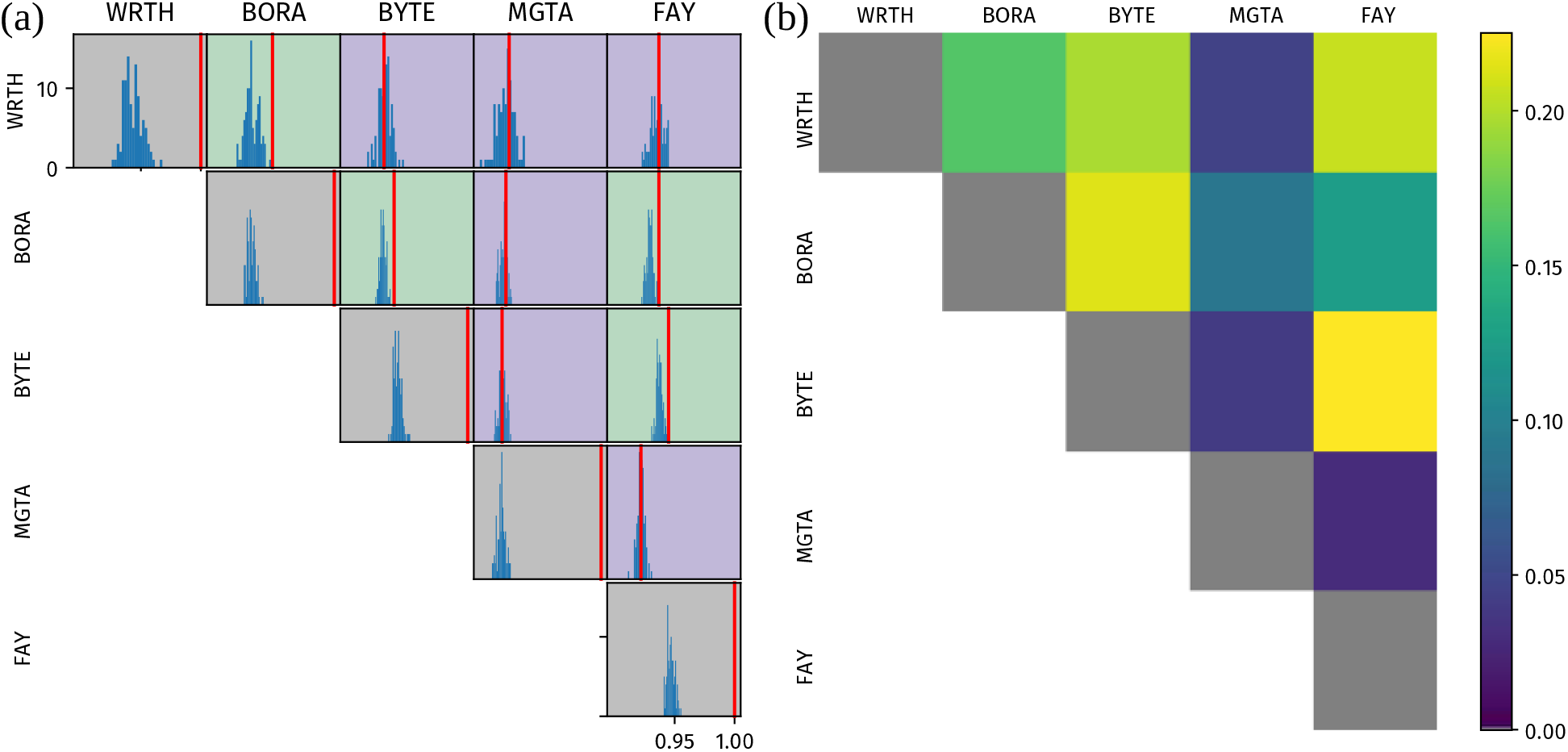
(a) Synchronisation of hourly-activity time-series between pairs of hyenas. *x*-axes in all subplots represent the synchronisation score (see Methods), and the *y*-axes show frequency. Since hyenas generally follow very similar activity patterns, *x*-axes in all cases are between 0.9 and 1.0. Red lines show the true level of synchronisation for each pair, and blue histograms show the level of synchronisation when the order of days is shuffled (permutation test). Pairs where the true level of synchronisation is greater than 95% of shuffled cases are shaded green, and pairs where it is not are shaded purple. Grey shading indicates self-pairs, which by definition are completely synchronised and are included here as a sanity check. 4 pairs of hyenas show a more extreme synchronisation score than expected based on permutations. In all cases, the true synchronisation is on the right-hand side of the corresponding shuffled-days distribution of synchronisation, indicating an overall tendency for greater synchrony rather than anti-synchrony as compared to the null model. (b) Proportion of time pairs of hyenas spent within 200 m of each other. Grey indicates self-pairs, as in (a).

## 4 Discussion

In this work, we developed a behavioural classifier that can reliably detect basic behavioural states of spotted hyenas from accelerometer data, and used it to characterise hyena activity patterns throughout the day at high temporal resolution. We found that the five female hyenas we studied did not compensate for high activity days on subsequent days. Furthermore, there was a statistically, but not biologically, significant effect of individual identity on activity patterns, suggesting that individuals do exhibit some distinctiveness in their activity patterns but that the variation within individuals is on par with the variation between individuals. When considering synchronisation across individuals, we found that some pairs of hyenas displayed activity patterns that were more synchronised than expected based purely on daily patterns. These specific pairs were also those that had spent more time in close proximity, however proximity did not guarantee activity synchrony.

Overall, our work highlights the promise of a classifier-based approach to behavioural recognition, though we note that care must be taken to avoid pitfalls. Here, we used a two-pronged approach to establish both the technical and contextual validity of our classifier. First, we performed audit-wise and individual-wise cross-validation, which avoided the spuriously high accuracy rates that can appear in non-independent and auto-correlated data such as behavioural sequences. We found that high accuracy can still be achieved even when training on different individuals than those whose behaviour is being predicted, demonstrating that the classifier can effectively generalise across individuals. Second, as a test of contextual validity, we compared the daily patterns predicted by our classifier to those reported previously based on direct observations [18]. Consistent with previous observations, we found a repeating 24-hour nocturnal activity pattern, with two peaks in activity, one following the evening twilight, and one just prior to the morning twilight. The predictions by our classifier also align well with states characterised by VeDBA alone.

Although we have shown that our classifier can produce reliable behavioural sequences, it also has limitations. First, while the classifier performs well, it is still error-prone, especially with respect to distinguishing between the two static behaviours *LYUP* and *STAND*. This confusion is, in retrospect, to be expected based on the similar positioning of hyenas’ necks while sitting or lying with head up, and the fact that both are stationary behavioural states. The classifier is also limited in terms of the behavioural repertoire it can detect: our ground-truth data were collected only during the day, during playback experiments, and in a context where hyenas were mostly alone. Furthermore, the five-behaviour ethogram we used is a simple reduced model of hyena behaviour, limiting our analyses to basic movement-centric behavioural states. We cannot yet address other more complex behaviours such as those involved in, e.g., social interactions or hunting. Importantly, behaviours not among the five in our ethogram will still be labelled with a state from our ethogram, thus we must be cautious in our interpretation of the predicted behavioural sequences. The best interpretation for these results assumes that our behavioural sequences are a simple, albeit useful, approximation of the animals’ real behavioural sequences. Another limitation is that we can only assess classifier performance during the day, since observing hyenas in the night is a daunting task even with night-vision equipment, as a consequence of which classifier performance during the night is un-quantified. Finally, an obvious limitation of our study is the small number of hyenas from which we have data. The small sample size at the level of individuals may limit the generality of our findings, and furthermore restricts the types of questions we are able to address. In addition, the limited duration of our sampling period also means that we cannot address questions pertaining to longer timescales, e.g., seasonal or phenological changes. Nevertheless, for each of the five individuals, our approach does enable us to obtain continuous behavioural records on a second-by-second basis, across an extended recording period of several weeks. These continuous behavioural records allow us to characterise hyena activity patterns and to address several relevant biological questions.

First, we found that hyenas in our study did not seem to make up for high activity days by being more inactive on subsequent days, or vice versa. Sleep, a closely related circadian rhythm, is found to be compensated in other species in what is known as sleep rebound [32]. While animals can make up for sleep debt straightforwardly by sleeping more hours [47–49], excess activity can also be compensated for in more complex ways. For instance, across several days, through physiological and behavioural changes, the total energy expenditure of primates of the same species does not scale with activity [50], so that animals which are differently active might be expending similar amounts of energy in a day [51–53]. Analogously, compensation could happen physiologically, by reducing energy output while not perceptibly reducing activity levels. Alternatively, straightforward activity compensation of the sort we describe might indeed be occurring, but at a time-scale different to a day. While we have shown that this does not happen across two- or even five-day timescales (Appendix C), hours of high activity could be compensated for in consecutive hours. Conversely, compensation could also occur on a much longer time-scale such as weeks or months. Activity compensation could also come into effect exclusively on days of exceptionally high activity. Longer-term biologging studies could address this possibility directly through slight modifications of our methods. Similar studies could also monitor various environmental and internal variables for individuals, controlling for which activity compensation can perhaps be detected. Perhaps the simplest explanation, of course, is that there is no activity compensation in this species.

Second, there was a statistically significant effect of the identity of an individual on its daily activity pattern. Within-individual variability in activity curves was slightly lower than across-individual variability. However, this effect is quite weak, and the statistical significance arises mainly due to the very large number of pairs of days. An alternative formulation of this analysis that is not affected by the large number of day-pairs (Appendix D) finds a similar result, with some support for individuality but not much statistical significance. While the influence of individual identity on circadian rhythms has been studied in humans [35], there are few analogous studies in animals (e.g., dominance hierarchy is known to influence activity patterns in rodents [2].) Our study uses naturally occurring activity patterns to examine the effect of individual identity on activity patterns. Individual identity might affect circadian rhythms through internal as well as social factors. In an animal with a complex social structure, where several social factors affect individual strategies, each individual’s activity pattern could be affected by various such social factors. For example, animals occupying different positions in the dominance hierarchy might be expected to show different activity patterns, or other factors such as age, sex, reproductive state, and prior experiences could also contribute to different activity patterns [54–56]. Due to the limited number of hyenas studied here we are currently unable to test whether these factors affect activity patterns, however using the same approach on a larger sample could allow us to determine what factors drive these differences. Finally, it is also possible that individual differences perceived here are caused by consistent external factors acting differently for different individuals, e.g., as a result of differing ranging patterns (which in itself would be individual variation). The variation in hyena activity curves, while not affected strongly by individual identity, is likely affected by various other factors.

Variability in hyena activity patterns may also emerge as a result of differing environmental and social conditions among these individuals. One simple manifestation of such a social interaction could occur as synchronisation of activity patterns [56]. Here we found that some pairs of hyenas, if they had spent substantial time in close proximity, had more synchronised behaviours than expected by chance. None of the pairs of hyenas had synchronisation *less than* expected by pure chance. Our permutation tests suggest that the synchronous patterns were not purely driven by similar daily patterns for individuals ranging in similar areas. It is, however, possible that the synchronisation we detect could arise from responses to the same local temporally varying environmental factors that affect specific pairs of hyenas. Such responses would be difficult to disentangle from direct social influences. While time spent together seemed to be a prerequisite for activity synchrony, the converse was not true. For example, two pairs (WRTH-BYTE and WRTH-FAY) spent more time in proximity than did the pair BORA-FAY; and yet the former pairs showed no evidence of activity synchronisation while the latter pair did. Our data thus suggest that proximity is a necessary, but not a sufficient, condition for synchronisation. Social interactions can entrain activity patterns leading to synchronisation [57–59], and proximity, which is a necessary condition for most social interactions, allows the potential to synchronise for each pair, but whether hyenas do or do not synchronise seems to depend on other factors. In contrast, a lack of proximity effectively guaranteed that synchronisation did not occur, at least at an aggregate scale. For example, the individual MGTA spent very little time in proximity to other study hyenas, and also did not synchronise with any of them. Since synchronisation as well as its absence are seen in hyenas naturally and without experimental intervention, this points to differential levels of synchronisation between hyenas in the population as a whole. Because of the limited number of individuals in this study, it was not possible to perform statistical tests to compare networks of proximity and synchrony, nor were we able to further investigate the potential drivers of synchrony vs lack of synchrony. Further studies on this and other species with a greater number of tagged individuals could overcome this issue. Additionally, we stress that we have compared *overall* synchronisation between activity patterns for a pair of hyenas with their *overall* proximity. This is different from comparing synchronisation when the individuals are in proximity against when they are not. Arguably, this is a stronger result—if we were to compare synchronisation between activity patterns only when a pair of hyenas were near each other, we expect that we would see high synchronisation values. This is because for a hyena pair to be in proximity, it is likely that the hyenas are in the same activity state (e.g., moving together, or both at rest). Future work using similar methods could address more specifically the proximate drivers of synchrony vs lack of synchrony, i.e. whether individuals synchronise their activities only in certain contexts. Finally, what social factors determine whether synchronisation occurs, and conversely, whether synchronisation enables or restricts some social interactions, also merits further study.

Overall, our work highlights the feasibility and value of accelerometer-based behavioural classification for studying animal activity patterns in the wild. While we focus here on spotted hyenas, the approach taken is not species-specific and we therefore expect it to be applicable to other study systems, opening up new avenues of exploration to understand the drivers of activity patterns across species.

## Ethics

All data were collected with the full authorisation of relevant Kenyan and U.S. institutions. These included license No. NACOSTI/P/22/19596 from the Kenyan National Commission for Science Technology and Innovation, capture permit No. KWS/904 from the Kenya Wildlife Service, and permit No. WRTI–188-05-22 from the Kenyan Wildlife Research and Training Institute. All research followed guidelines of the American Society of Mammalogists [60], and was also approved by the IACUC at Michigan State University as protocol No. PROTO202200047.

## Data and code availability

Raw GPS and accelerometer data are publicly available [61]. Code for behaviour classification, as well as for biological analyses, are available on github at https://github.com/pminasandra/hyena-acc and as a stable upload associated with this manuscript on Zenodo [62].

## Authors’ contributions

The hyena collaring study was conceived by ASG, FHJ, ASP, and KEH, who also secured funding and collected the bio-logging data used in this work. KEH maintains the long-term field site of spotted hyenas in Masai Mara National Reserve, Kenya. FHJ and KEH tested the collars. ASG, FHJ, ASP, and KEH collected the data. PM developed and validated the classifier, performed the biological analyses, and wrote the initial manuscript with inputs from ASP. EDS performed a code review. All authors contributed to, and approved, the final manuscript.

## Acknowledgements

We thank the Kenya Wildlife Service, the Kenyan Wildlife and Research Training Institute, the Kenyan National Commission on Science, Technology and Innovation, and the Narok County Government for permission to conduct this research. We thank Benson Pion, Rebecca LaFleur, and Morgan Lucot for field assistance, and Max Martin and Kenna Lehmann for annotating video files for ground truth. We thank Hester Brønnvik, Laura Smale, and Marie Roch for valuable comments on the methods and the manuscript. We thank Mark P Johnson for providing the DTAG technology used in the collars. We thank Reviewers 1 and 2 for suggesting the analyses in Appendices C-E. We also thank Reviewer 1 for referring us to useful literature.

## APPENDIX A Classifier performance

The Random-Forest classifier was the most accurate (92%), followed by the k-NN classifier (88%) and the SVM classifier (87%). The precision and recall scores for each of these classifiers, in the same order, are provided in Table A1.

We also show the confusion matrices in different testing paradigms (see Methods) for the *k*-NN and SVM classifiers here (Figure A1, Figure A2).

**Table A1:** Precision and recall values for each behavioural state for each classifier. The *LYUP* -*STAND* confusion is more apparent with these numbers.

| Random Forest |  |  |  |  |  |
| --- | --- | --- | --- | --- | --- |
| - | LYING | LYUP | STAND | WALK | LOPE |
| Precision | 1.00 | 0.84 | 0.72 | 0.96 | 0.93 |
| Recall | 0.99 | 0.70 | 0.80 | 0.97 | 0.98 |
| *k*-Nearest-Neighbours |  |  |  |  |  |
| - | LYING | LYUP | STAND | WALK | LOPE |
| Precision | 1.00 | 0.69 | 0.71 | 0.92 | 0.91 |
| Recall | 0.99 | 0.73 | 0.58 | 0.96 | 0.98 |
| Support Vector Machine |  |  |  |  |  |
| - | LYING | LYUP | STAND | WALK | LOPE |
| Precision | 0.99 | 0.72 | 0.61 | 0.91 | 0.91 |
| Recall | 0.99 | 0.52 | 0.64 | 0.96 | 0.98 |

**Figure A1:**
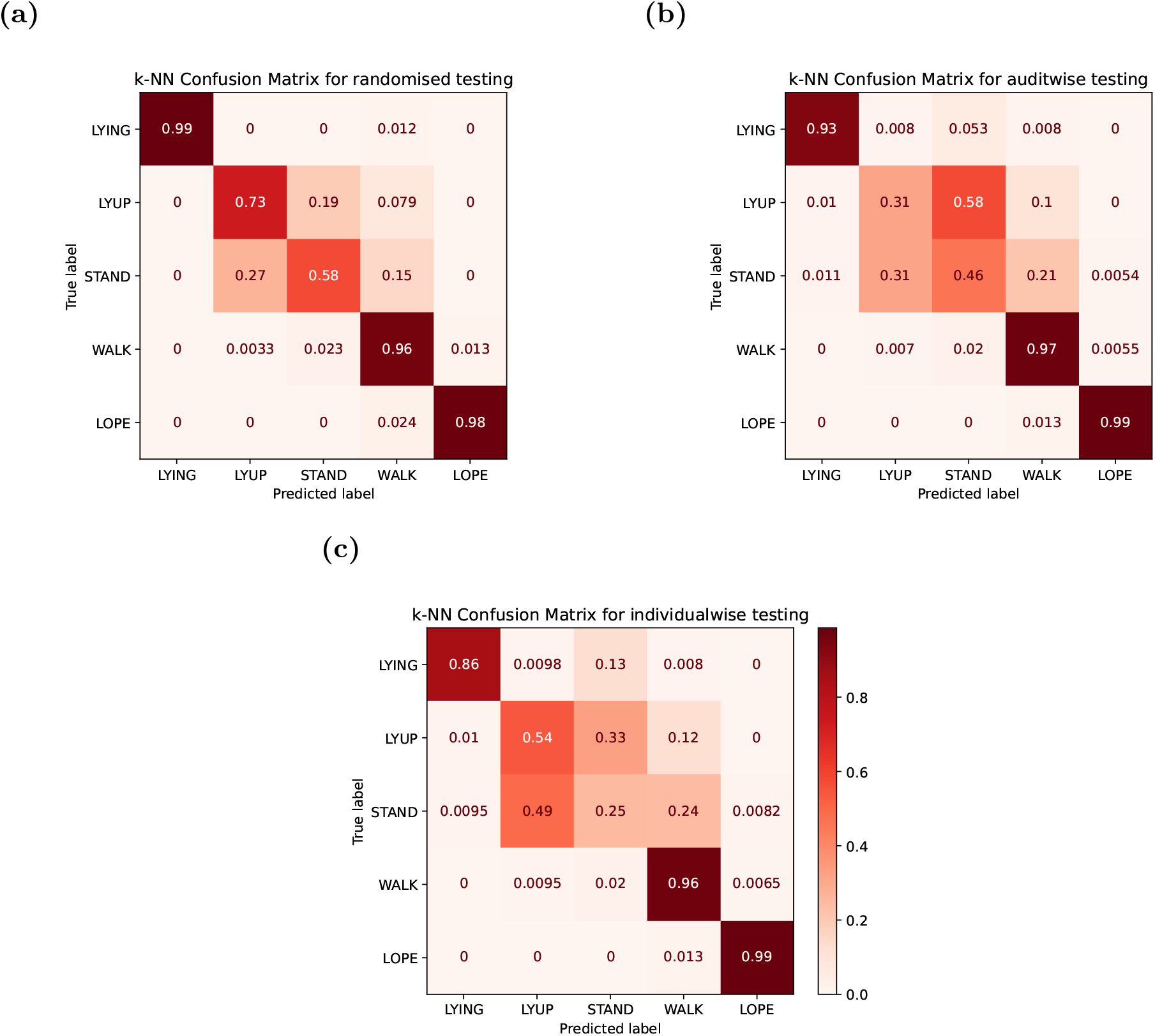
Confusion matrices for *k*-Nearest Neighbours classifier

**Figure A2:**
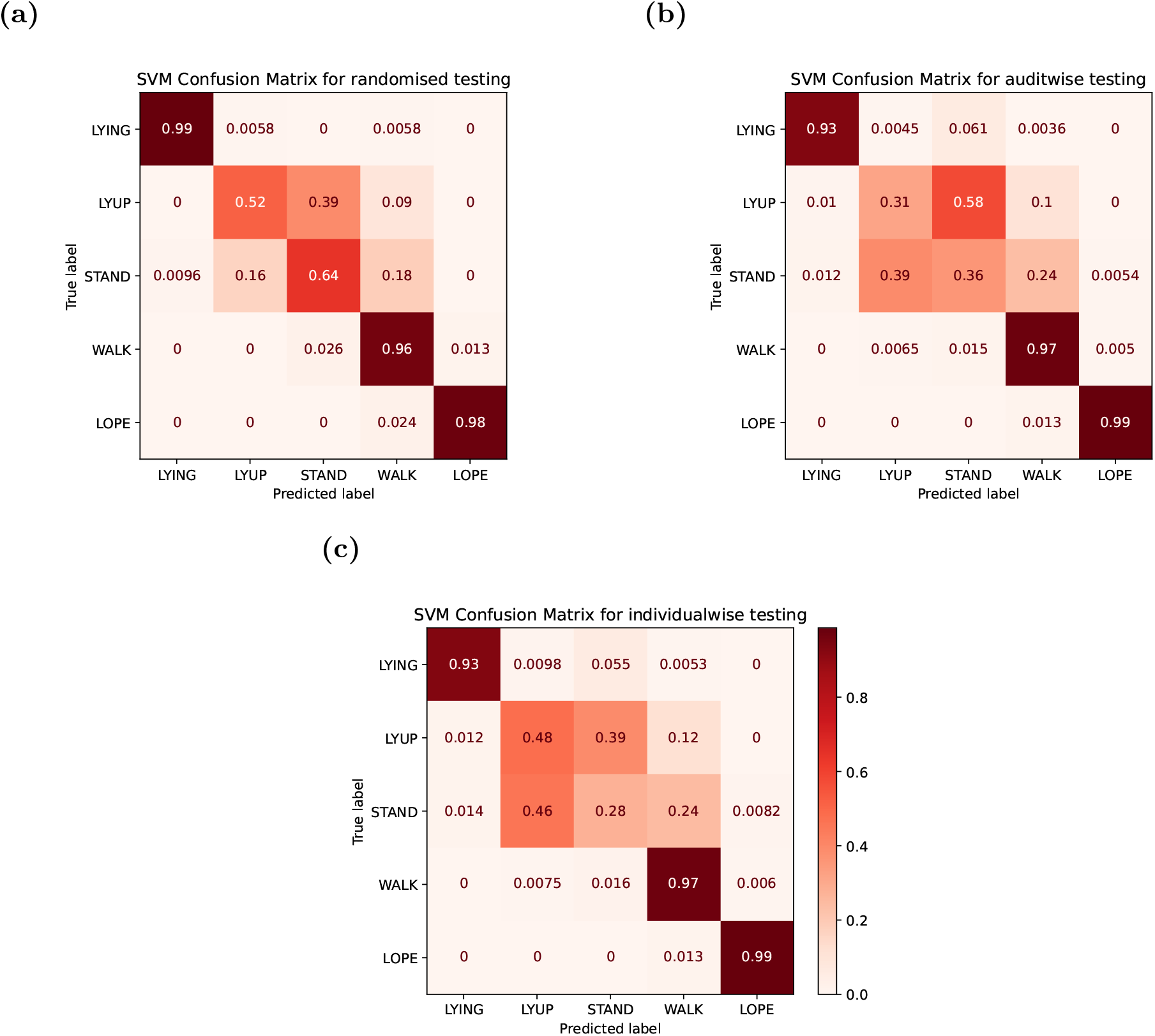
Confusion matrices for a Support Vector Machine classifier

## APPENDIX B Definitions of permutation tests

### B.1 Quantifying individual idiosyncrasies

This section describes how we computed the average difference between within-individual and across-individual variability of activity patterns between days. Let *d*_*A,j*;*B,k*_ be the distance between activity patterns of hyena *A* on day *j* and hyena *B* on day *k*. We defined

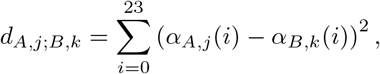

where *i* is an hour of the day between 0 and 23, and *α* is the proportion of time spent in an active behavioural state.

First, we computed the average within-individual distance between all pairs of days for each hyena,

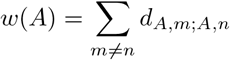

and then computed the average of these value across all hyenas to obtain *w*. We then computed the across-individual distance between all pairs of days for each unique pair of hyenas,

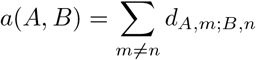

and, as before, we computed the average of these values across all pairs of hyenas to obtain *a*.

The variable *a − w*, the difference between the mean across-individual and mean within-individual distances between activity patterns was chosen as a test statistic.

### B.2 Quantifying synchronisation of activity patterns

The synchronisation score for activity patterns was defined as

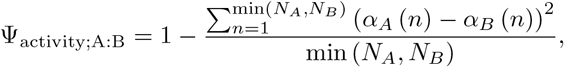

where *N*_*A*_ and *N*_*B*_ are the total number of hours for which data was available for hyenas A and B, and *α*_*A*_(*n*) and *α*_*B*_(*n*) are the proportions of time spent by A and B respectively in active behavioural states. This metric becomes 1 when the activity patterns are perfectly identical, and can theoretically attain a minimum value of 0.

## APPENDIX C Activity compensation across several days

Activity compensation might not occur from day to day. To test if consistent high or low activity across several multiple days was compensated for, we calculated the average activity on all *m*-consecutive-day tuples, and used them to predict the activity level on the following day using a linear regression. We did this for *m* = 2 and 5.

Figure C1 shows that no clear trend was seen in activity levels across days, even when accounting for activity across several days.

**Figure C1:**
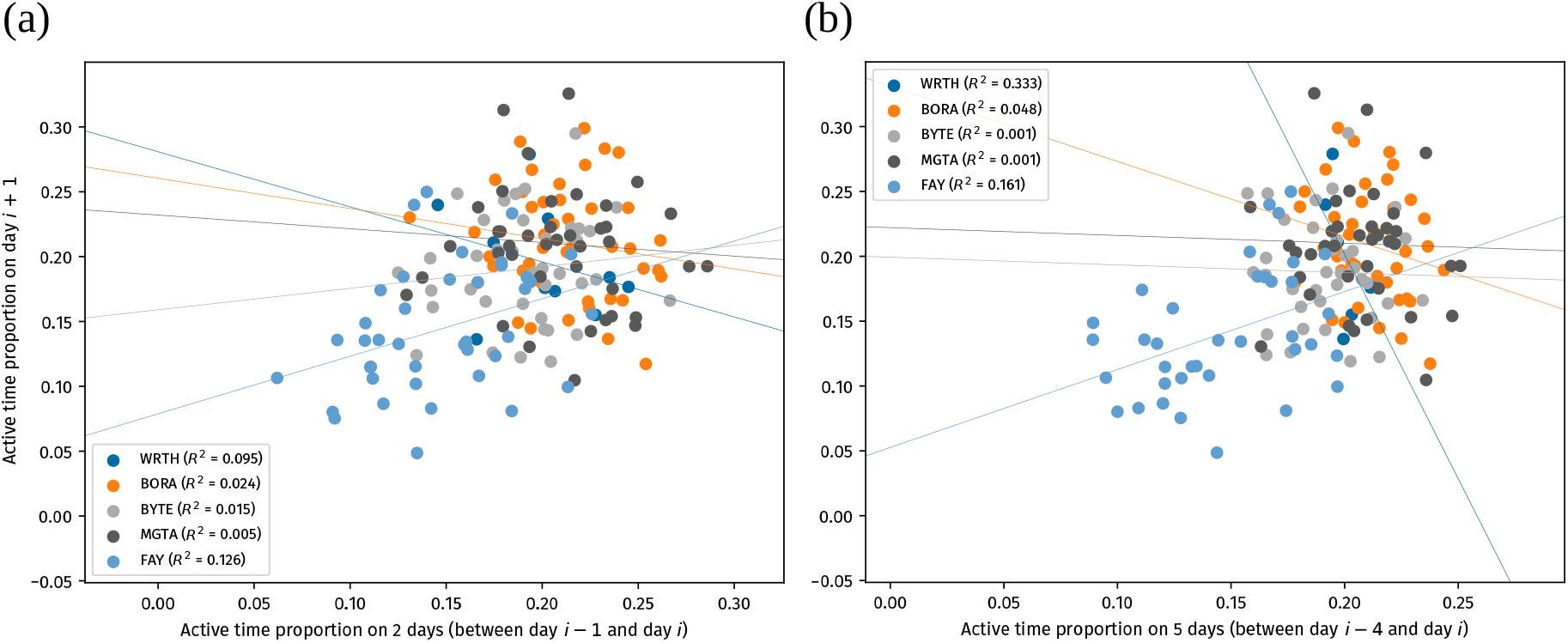
No compensation of activity, or other general trend, is seen when accounting for average activity over the preceding (a) 2 days or (b) 5 days.

## APPENDIX D Alternate method for quantifying individual variability

In the main text, we found a statistically significant effect of individual identity, but with a negligibly small effect size. This happens because of the very large number of day-pairs that results from several days of data across five hyenas. Here, we provide an alternate formulation of the analysis that overcomes this issue. To quantify the day-to-day variability of an individual hyena, we define a statistic as follows. First, for each hour of the day *i*, we find the across-days variance in the activity level (see Methods), *v*_*i*_. We then find the sum of these hourly variances in activity, the total daily variability of the individual *j*’s activity pattern, as

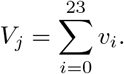

We then calculate a similar daily variability statistic while ignoring individual identity. To do this, we take all activity curves available, from all hyenas, in a pool, and randomly assemble a sequence of days drawn from all individuals, with this sequence being of equal length as the observed data for individual *j*. We repeat this 5000 times, and each time, we compute the above metric for the resampled data-set. This provides a null model of the variability in the individual’s activity pattern.

The effect considered is the difference between the mean of the resampled data-set, 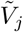 and the variability observed for the individual,

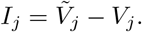

Positive values of *I*_*j*_ indicate that the individual’s activity pattern is less variable than the variation of activity curves across all individuals. Negative values indicate that the individual’s activity pattern is, conversely, more variable across days.

We calculated the metric *I*_*j*_ for all individuals, and computed *p*-values based on their definition, assuming a 2-tailed *p*-value. Likewise, we found the overall individuality score *I*, and computed the *p*-value there similarly. We found that, on average, individuals are slightly less variable than expected from the permutations, although this result was not statistically significant. (*I* = 0 .071; a nd *p* = 0.455) (Figure D1).

Overall Individuality scores *I*_*j*_ and *p*-values for our individual hyenas are: *I*_WRTH_ = 0.135, *p* = 0.05; *I*_BORA_ = 0.018, *p* = 0.323; *I*_BYTE_ = 0.145, *p <* 0.0002; *I*_MGTA_ =*−*0.144, *p* = 0.0002; and *I*_FAY_ = 0.208, *p <* 0.0002. MGTA is more variable across days than the overall variation across the group, while other individuals are more idiosyncratic.

**Figure D1:**
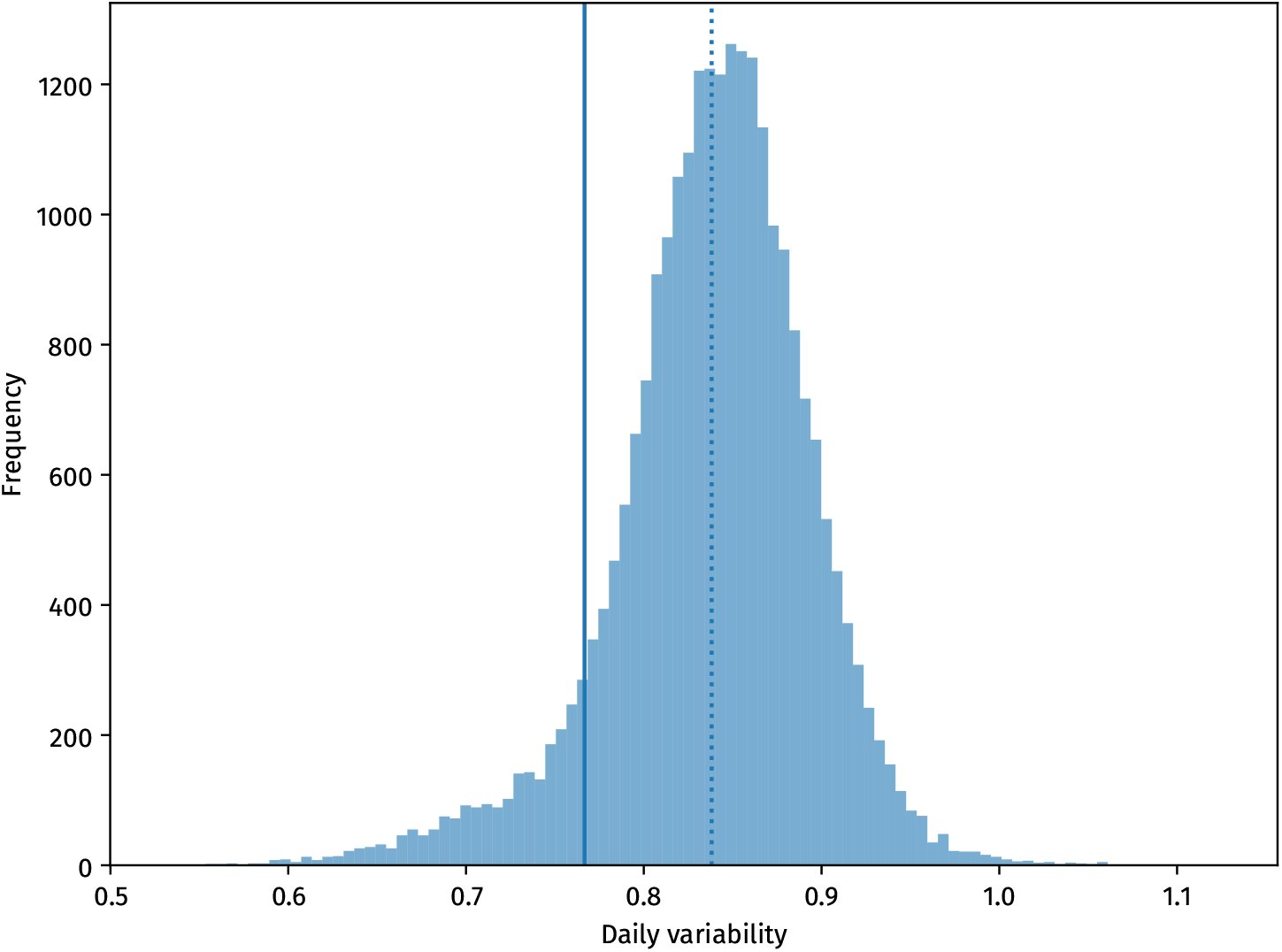
Histogram of the daily variabilities of permuted data-sets, i.e., variability across individuals (translucent bars, with dotted line indicating the mean); and actually observed average daily variability (bold vertical line).

**Figure E1:**
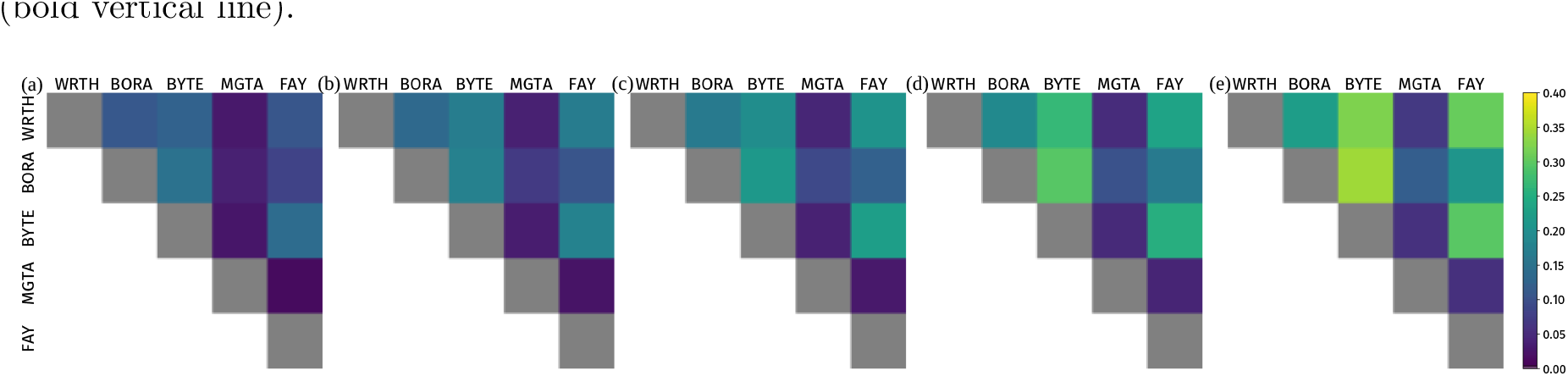
Proximity networks constructed with proximity threshold distances (a) 50 m, (b) 100 m, (c) 200 m, (d) 300 m, and (e) 500 m. Proportion of time spent in proximity always increases with increasing threshold for every pair, as expected.

## APPENDIX E Proximity network with varying thresholds

We constructed proximity networks (see Methods) with five different threshold distances, 50 m, 100 m, 200 m, 300 m, and 500 m, as a test of consistency. We found that the networks showed consistent ordering of proximity for all pairs (Figure E1). As we increased the threshold distance, the proportion of time spent in proximity for a pair always increases. This is sensible because, e.g., if a pair of hyenas spends 10% of their time less than 50 m from each other, the proportion of time spent less than 100 m away has to be *at least* 10%.

Since our results compare the proportion of time spent in proximity across different pairs, our results are not affected much by choosing a specific threshold.

